# Feeder-Free Generation of Lymphatic Endothelial Cells from Human Induced Pluripotent Stem Cells

**DOI:** 10.64898/2026.03.19.712968

**Authors:** Aditi Prasad, Shrey Patel, Simon Ng, Clifford Z. Liu, Bruce D. Gelb

## Abstract

**Abstract:** The lymphatic system is essential for maintaining fluid homeostasis, lipid transport and supporting immune function. Despite its central role in health and disease, advancements in understanding human lymphatic vasculature has been constrained, in part because primary human LECs are difficult to access and study in disease-relevant contexts. This study describes an efficient and scalable feeder-free method to differentiate human iPSCs into lymphatic endothelial cells (LECs) that are transcriptionally and phenotypically similar to primary fetal LECs. An iPSC-derived LEC system overcomes a drawback of primary cells by enabling precise genetic perturbations, supporting study of lymphatic diseases of interest in a human context. By grounding our approach in *in vivo* stages of lymphangiogenisis, we describe a staged protocol that recapitulates the key milestones of lymphatic development. We first adapted a published method to differentiate human iPSCs into venous endothelial cells (VECs) and then initiate transdifferentiation of VECs into LECs. Using immunocytochemistry, qPCR, as well as flow cytometry, we demonstrated expression of lymphatic-specific markers in the differentiated population. We further characterized our induced VECs (iVECs) and LECs (iLECs) through bulk RNA sequencing analysis and compared the populations to pseudobulk VEC and LEC transcriptomic datasets generated from human fetal heart endothelia at 12, 13 and 14 weeks of gestation. Through this work, we expanded the repertoire of approaches for accessing LECs, with the goal of accelerating discoveries in lymphatic biology and therapeutics.

**Abstract summary image:** 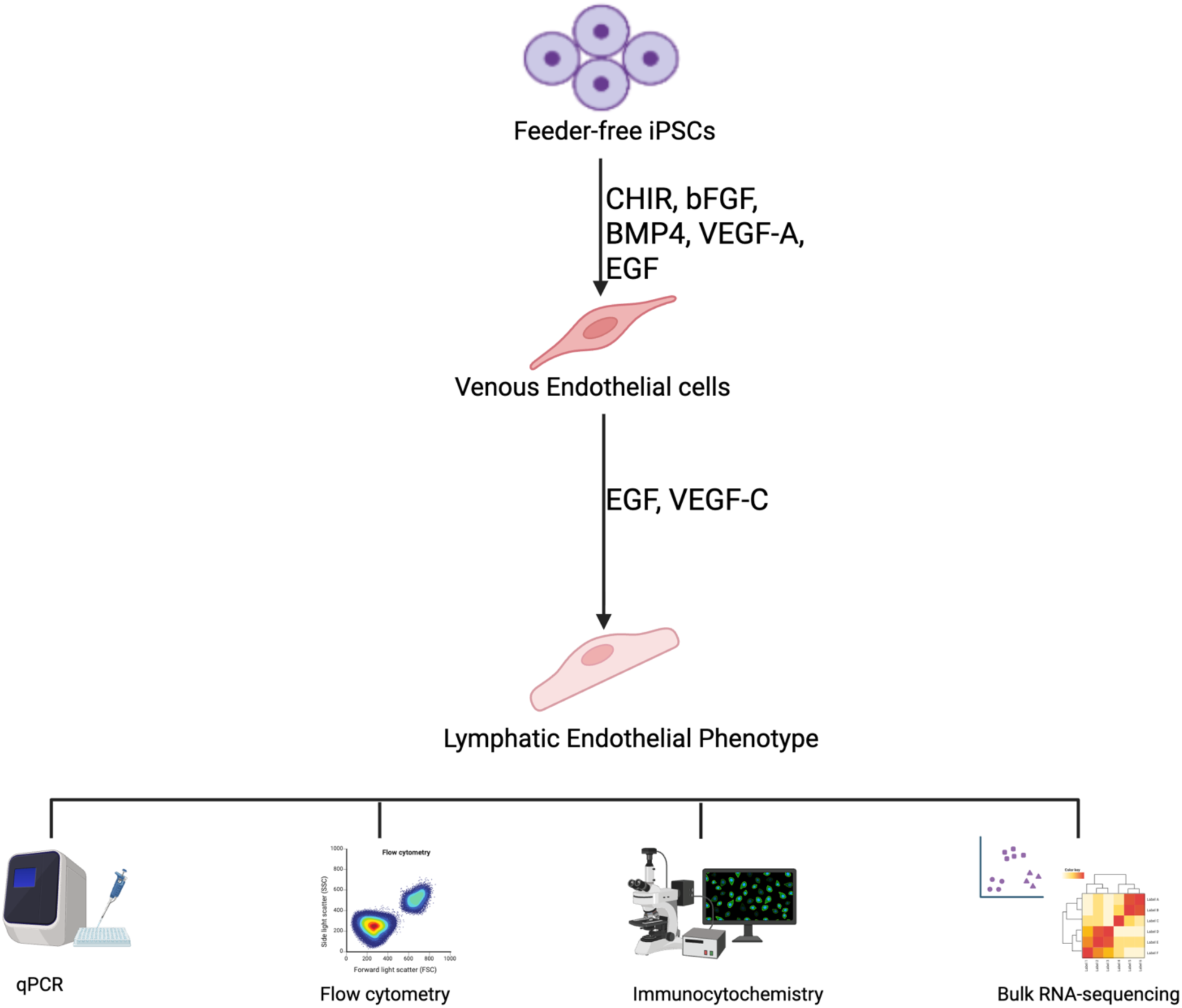

## INTRODUCTION

The lymphatic system is a network of capillaries within interstitial spaces of cells, which connect back to larger lymphatic vessels called collecting lymphatics, where tissue fluid is drained and eventually transported back into the blood circulatory system. All lymphatic vessels and capillaries consist of lymphatic endothelial cells (LECs), which align to form into tubes. The lymphatic system plays a pivotal role in maintaining tissue and fluid homeostasis, immune surveillance, as well as facilitating absorption of lipids into the blood.^1^ Impairments in the development and formation of lymphatic vasculature can lead to a myriad of pathologies, including lymphedema and lymphangiectasia, and can contribute to cancer metastasis.^2–5^ Despite the importance of the lymphatic vasculature in maintaining fluid homeostasis and the detrimental effects of lymphatic disease, cellular mechanisms that drive lymphangiogenesis still remain incompletely understood.

Prior work has guided our understanding of lymphatic development *in vivo*, showing that the mammalian lymphatic system develops sequentially during fetal life and is moderated by a host of key regulatory molecules that drive a subpopulation of cells in the cardinal vein to bud off from the dorsal cardinal vein to form primitive LECs, which eventually develop into the lymphatic vasculature. Mature LECs express *PROX-1*, a master lymphatic regulator, along with *LYVE-1*, *PDPN*, *SOX-18*, *COUP TF-II* as well as *FLT4*, which encodes VEGFR3, the receptor for the VEGF-C ligand.

Developmental models are a crucial tool to elucidate mechanisms that drive the developmental trajectory of the lymphatic system, and recent work in the field has described various *in vitro* and *in vivo* systems that have helped guide our understanding of lymphangiogenesis and lymphatic disease. Mouse models *in vivo, ex vivo*, and *in vitro* have been one of the commonest systems used to study lymphatic development. The use of purified *ex vivo* embryonic mouse LECs in culture has contributed to understanding the pro-proliferative role played by Fgfr1 and its ligand Fgf2 during lymphangiogenesis as well as the cooperative role of Fgfr2 signaling with Vegfr3 signaling in lymphatic tube formation.^6^ Mouse embryonic stem cells (mESCs) grown in suspension (embryoid bodies) have also been shown to spontaneously differentiate and adopt a lymphatic-like phenotype by Day 18 of culture.^7,8^ The VEGFR3/VEGF-C signaling pathway has been emphasized as a crucial signaling pathway for LEC development.^9–11^

Although *in vivo* animal models provide a robust system to study development and some diseases of the lymphatic vasculature, a major drawback of such models is that they can have low translationability of therapeutic findings into humans.^12^ The use of primary human dermal LECs has allowed researchers to uncover mechanisms of lymphatic proliferation, tube formation and migration in an *in vitro* system.^13,14^ Another approach is to differentiate human induced pluripotent stem cells (iPSCs) into LECs. However, all published protocols for doing so require the use of some form of feeder cells–mouse embryonic fibroblasts or OP9 stromal feeder cells.^15,16^ The use of feeder cells in a protocol has various limitations: technical hurdles, impact on reproducibility, increased costs, and is time consuming. Additionally feeder cells secrete cytokines, which may engage in paracrine signaling with the differentiating cells, preventing complete control of the cytokines with which the differentiating cells interact.^17,18^

Our study aimed to elucidate a method to efficiently differentiate human iPSCs into LECs in a feeder-free, serum-free condition and, in so doing, circumvent the limitations of a feeder system. Given the prior knowledge of LECs’ trans-differentiation from a venous endothelial cell (VEC) population, we chose to first efficiently differentiate iPSC into induced VECs (iVECs)^19^, from which we then drove transdifferentiation into induced LECs (iLECs). With the understanding that EGF and VEGFR3/VEGF-C signaling is important for LEC fate determination, we included these cytokines in our lymphatic differentiation conditions.

## RESULTS

### Monolayer differentiation of feeder-free iPSCs into a robust endothelial cell progenitor population

Endothelial cell lineages are known to derive from mesoderm during embryogenesis, and establishment of a robust endothelial population is crucial to further vascular lineage development.^20,21^ Recent literature has pointed to the use of the GSK3 inhibitor, CHIR, in efficiently differentiating iPSCs into mesoderm derivatives in monolayer, feeder-free conditions.^22,23^ In order to differentiate human iPSCs into an endothelial progenitor (EP) population, we adapted a previously established protocol, which utilized CHIR as well as VEGF and BMP4.^19^ WTC11 cells were plated as a monolayer and induced to form a primitive streak using CHIR, followed by mesoderm differentiation using bFGF. FGF signaling is well established as a key regulator of epithelial-to-mesenchymal transition and of lateral and paraxial mesodermal fate specification at the primitive streak.^19,23–25^ VEGF and BMP4 were then introduced to push the monolayer towards EP fate over a 48-h period (**Figure 1A)**.

**Figure 1.**
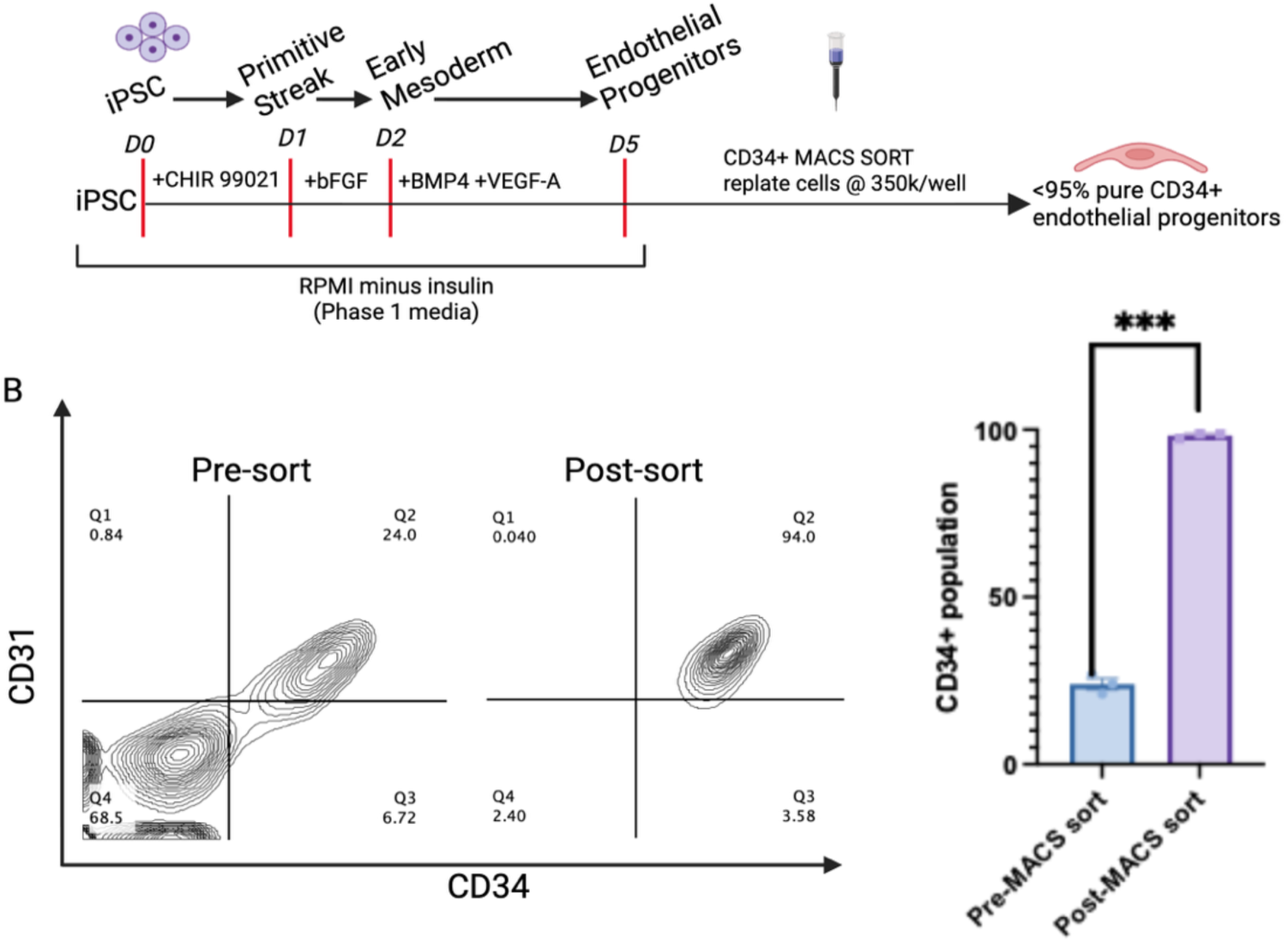
Differentiating iPSCs into a pure endothelial cell progenitor population in a monolayer feeder-free condition. **A.** Schematic representation of iPSC differentiation into EPs. **B.** Flow cytometry plot representative of pre and post CD34 sorted Day 5 endothelial progenitor cells. Cells were labeled with CD31 and CD34 endothelial markers. CD34+ cell efficacy quantified pre- and post-MACS sort (n = 3, ***p < 0.001, 2-way ANOVA, Šídák’s multiple comparisons test)

To purify the EP population, we employed magnetic-activated cell sorting (MACS), to select for a CD34+ population. This technique allowed us to routinely purify our EP progenitor population to a 95% pure population. To determine the efficiency of the cell sort, flow cytometry was used to analyze the pre- and post-sort populations for CD31^+^/CD34^+^ cells. The efficiency of the WTC11 line in differentiating into the CD34^+^/CD31^+^ population was ∼23 ± 2%, and the overall sorting efficiency was over ∼98 ± 0.5% pure for CD34^+^ cells in the post-sort analysis. Additionally, ∼94% of the WTC11 sorted cells were CD34^+^/CD31^+^. Subsequently, the purified EPs were re-plated on Matrigel-coated 24-well plates and were treated to adopt an iVEC fate (**Figure 1B**).

### EGF and bFGF drive differentiation of a robust venous endothelial cell population from a pure endothelial progenitor population

Preliminary work demonstrated that VEGF-C/VEGFR3 signaling alone is insufficient to drive the direct differentiation of EPs into LECs (data not shown). To achieve our goal of generating iLECs, we sought to recapitulate key stages of *in vivo* lymphatic development within our *in vitro* system. In view of the understanding that LECs transdifferentiate out of the cardinal vein *in vivo*, it was imperative to first create a robust iVEC population, which could then be pushed towards a lymphatic fate. To differentiate iVECs, we adapted a previously published eight-day protocol that used a monolayer feeder-free approach (**Figure 2A**).^19^

**Figure 2.**
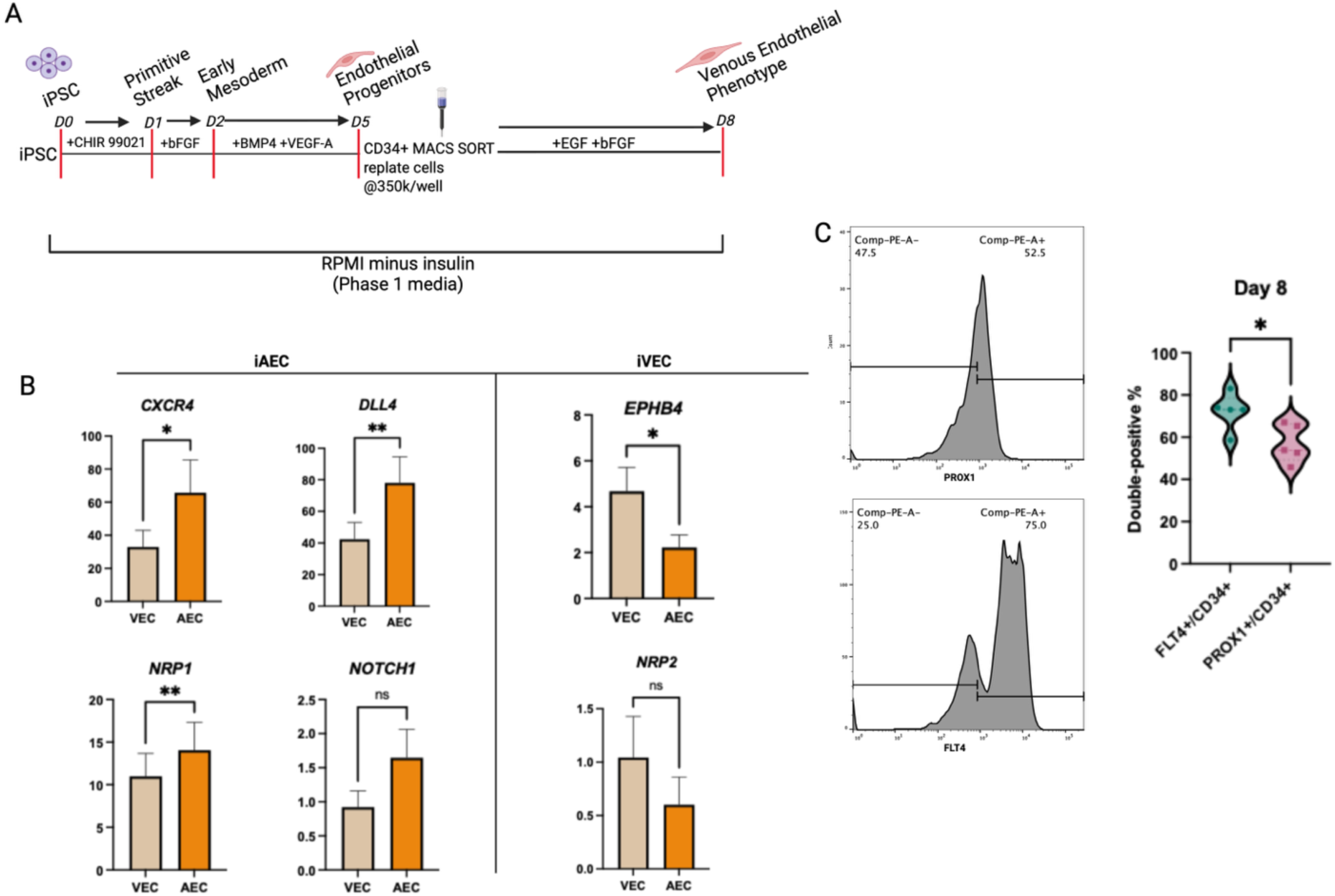
EGF and bFGF drive robust differentiation of iPSCs to venous endothelial cells. **A.** Schematic representation of an eight-day protocol to differentiate iPSCs into induced venous endothelial cells. **B.** qRT-PCR analysis on Day 8 shows upregulation of venous markers and downregulation of arterial markers in iVECs, iAECs act as a negative control (n=7, *p<0.5, **p<0.01, paired t-test). **C.** Flow cytometry analysis shows robust expression of FLT4 (75%) and PROX1 (55%) in the iVECs. iVECs expressing FLT4+/CD34+ were significantly higher in number than those expressing PROX1+/CD34+ (n=5, *p<0.5, paired t-test).

Purified EPs on Day 5 were treated with EGF (10 ng/ml) and bFGF (20 ng/ml) for 72 h to induce a VEC fate. Additionally, a small portion of purified EPs were treated with VEGFA (10 ng/ml) along with EGF (10 ng/ml) and bFGF (20 ng/ml) for 72 h and cultured separately to produce induced arterial endothelial cells (iAECs). iAECs were used as a negative control. qRT-PCR analysis of the Day 8 iVECs showed an upregulation of the venous gene markers *EPHB4* and *NRP2* and a downregulation of the arterial markers *CXCR4, DLL4, NOTCH1* and *NRP1.* Conversely, the iAECs showed an upregulation in arterial gene expression and downregulation in venous gene expression. Although differences in iVEC and iAEC expression levels for *NRP2* and *NOTCH1* were not significant, the expression patterns trended in the expected direction: higher *NRP2* and lower *NOTCH1* expression in iVECs (**Figure 2B**). Flow cytometry analysis of the iVECs for FLT4 and PROX1 expression revealed ∼75% of the population was FLT4+ and ∼55% was PROX1+ (**Figure 2C**). Expression of FLT4 was significantly higher than PROX1. *FLT4*, which encodes VEGFR3, is required to interact with the VEGF-C ligand. Upregulation of FLT4 demonstrated that the iVECs were primed to interact with VEGF-C and further transdifferentiate into an LEC fate. Moreover, a subpopulation of VECs also express PROX1 *in vivo*, marking these cells as the ones that will transdifferentiate out of the cardinal vein into the lymphatic sacs. It was, therefore, expected that the majority of the iVECs would express FLT4 significantly more than PROX. Overall, these results demonstrated differentiation of a robust venous endothelial cell population.

### EGF and VEGF-C promote transdifferentiation of iPSC-derived venous endothelial cells into iPSC-derived lymphatic endothelial cells

Following successful generation of iVECs, we sought to transdifferentiate them into iLECs, mirroring the *in vivo* process in which of LECs emerge from the cardinal vein. To do this, iVECs were dissociated and replated on Matrigel-coated 24-well plates at roughly 200k/well in Phase 2 media supplemented with EGF (10 ng/ml) and VEGF-C (50 ng/ml) for 10 days (**Figure 3A**). Flow cytometry analysis revealed robust expression of PROX1 (the master regulator of lymphatic development) and PDPN (a key marker of LEC maturity) in the differentiating cells. The proportion of PROX1⁺ iLECs was high at both time points, with 88 ± 4% at Day 14 and 94 ± 2% at Day 18 (mean ± SEM, n = 5 per time point). When combined, the overall PROX1⁺ population averaged 90 ± 2%. The proportion of PDPN^+^ iLECs was similar but slightly lower than the PROX1^+^ population with a 93 ± 3 % PDPN^+^ population on Day 14 and a 92 ± 4 % PDPN^+^ on Day 18. Additionally, the percentages of PROX1^+^ and PDPN^+^ populations of iLECs were not significantly different from Day 14 to 18, indicating that the differentiated cells were maintaining their lymphatic phenotype (**Figure 3B**, n = 5). Notably, flow cytometry density plots also revealed a small population (< 20%) of PDPN^+^/CD34^-^ cells. Mature LECs are known to lose CD34 expression while increasing PDPN expression. This population might reflect heterogeneity in the maturity stages of the iLEC population^26^. These data indicated that while most of the iLEC population might be fetal-like, expressing both CD34 and PDPN, a subset of the population might be losing CD34 expression as it moves down the development trajectory. Further, qRT-PCR analysis revealed robust expression of key lymphatic markers *NR2F2, FLT4, LYVE1, PROX1* and *SOX18* as well as the key endothelial marker *CD34*. Expression of all markers was maintained through Day 18 with no significant changes in expression levels (**Figure 3C**, n = 7). Taken together, these results indicated that treatment with EGF and VEGF-C was successful in differentiating cells that adopted a lymphatic phenotype and robustly expressed lymphatic markers.

**Figure 3.**
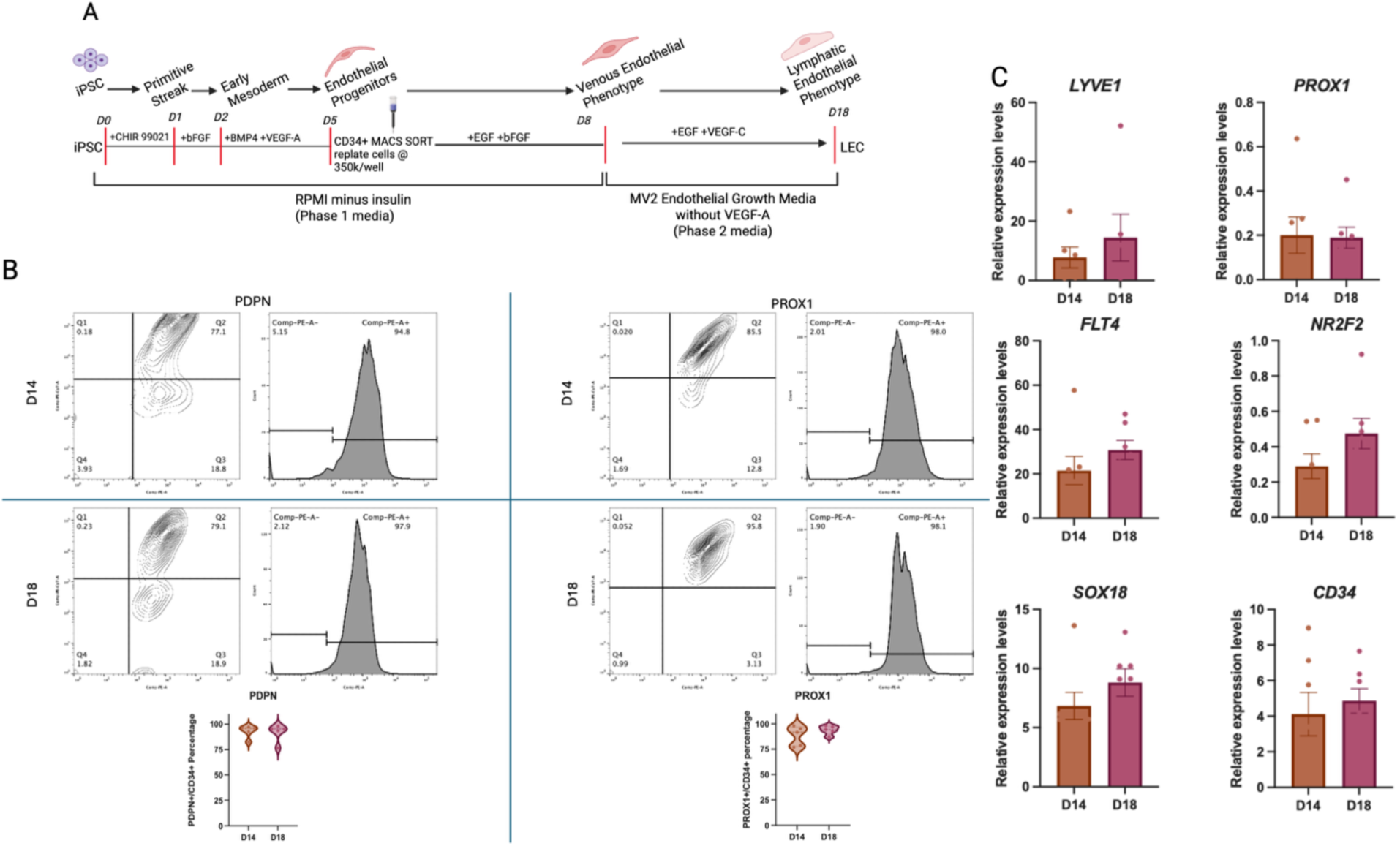
Day 18 analysis indicates transdifferentiation of venous endothelial cells into lymphatic endothelial cells. **A.** Schematic representation of the differentiation protocol using EGF and VEGF-C over a 10-day period to transdifferentiate iVECs into LEC phenotype. **B.** Flow cytometry analysis on Day 14 and 18 show robust expression of PDPN and PROX1 with no significant changes in the expression of these proteins between both time points (n=5, paired t-test). **C.** qRT-PCR analysis indicates strong expression of LEC and endothelial gene markers on both Day 14 (6 days post induction of LEC media) and Day 18 (10 days post induction of LEC media) (n=7, paired t-test).

### iLECs exhibit functional similarities to human dermal lymphatic endothelial cells

*In vitro* network and tube formation ability is an established standard method of characterizing angiogenic capacity of endothelial cells.^27–29^ In order to demonstrate that the iLECs preserved this function as well as expressed lymphatic marker proteins, we plated iLECs in 2D as well as carried out a tube formation assay, and then imaged iLECs in both environments with confocal microscopy. For the tube formation assay, we relied on previously published protocols and plated the iLECs in growth factor-reduced Matrigel with a 1.5-mm thickness on 8-well glass bottom plates. Within 4 h of plating, the cells rearranged into tube-like structures. iLECs robustly expressed both lymphatic- and endothelial-specific proteins when cultured in 2D as well as in tubes (**Figure 4A**). Differentiated cells showed marked expression of PDPN, PROX1 and LYVE1 as well as CD31, a pan-endothelial marker that is biased towards LECs.

**Figure 4.**
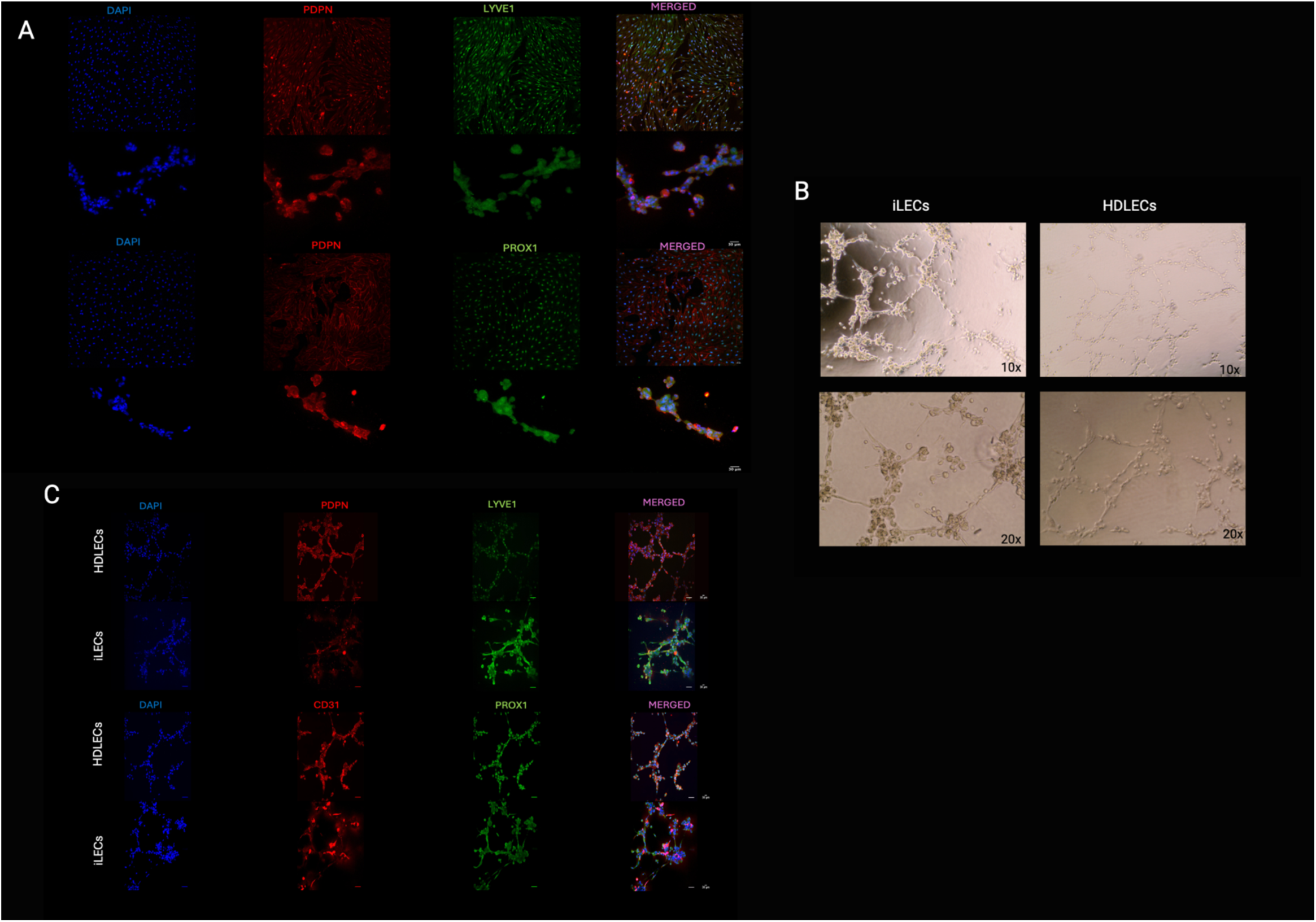
iLECs exhibit similar expression and functional properties to HDLECs. **A.** Confocal image of iLECs expressing PDPN, PROX1, LYVE1 and CD34 in monolayer and tubes (20x, Zeiss LSM880). **B.** Bright-field images of tube-like structures form by iLEC sand HDLECs (10x and 20x). **C.** Tube-formation assay indicating angiogenic capacity of iLECs similar to HDLECs.

Additionally, to assess if structures formed by the iLECs were in fact functionally relevant, we carried out the same assay and staining with primary human dermal LECs (HDLECs) as a positive control. The iLECs formed tube-like structures similar to the HDLECs (**Figure 4B**). Further, confocal microscopy images showed expression of lymphatic-specific proteins in the iLEC tube-like structures similar to the HDLEC tube-like structures (**Figure 4C)**. iLECs maintained expression of key lymphatic markers when they self-arranged into tube-like structures. Overall, these data indicated that the iLECs exhibited functional similarities and protein expression profiles when compared to primary HDLECs.

### iVECs and iLECs indicate similar transcriptomic profiles to fetal VECs and fetal LECs respectively

To determine the identity of our iVECs and iLECs, we sought to compare their transcriptomic expression profiles to those of fetal VECs and fetal LECs. To do this, we performed bulk RNA sequencing analysis of our iVECs and iLECs. In addition, we generated pseudobulk RNA sequencing datasets for fetal VECs and LECs using published single-cell RNA-seq datasets.

To conduct the bulk sequencing analysis, we collected cell lysates of iVECs on Day 8 and iLECs on Days 14 and 18. Leveraging packages in R, we normalized and variance-stabilized the data using DESeq2 and batch-corrected count read matrices using limma.^30,31^ As an initial step, we generated a PCA plot comparing our Day 8 iVECs to our Day 14 and 18 iLECs, as well as a gene set enrichment analysis (GSEA)(**Figure S1**). The PCA showed that the Day 8 iVECs clustered separately from the Day14 and 18 iLECs (**Figure S1A**). Furthermore, the GSEA heatmap revealed gene sets derived from fetal LECs were strongly enriched in both Day 14 and Day 18 iLECs compared with the iVECs, while sets such as “venous identity” and “venous origins” were enriched in our iVECs but not in the iLECs (**Figure S1B**).

Next, we compared our iVEC and iLEC transcriptomic data to previously published single cell RNA-seq datasets that sequenced human fetal heart endothelium at 13 and 14 weeks of gestation^32^ and a human fetal heart at gestational 12 weeks of gestation.^33^ We created pseudobulked expression profiles of our Day14 and 18 iLECs by averaging raw counts of both time points to create a singular iLEC variable. We then assessed the concordance between our iLEC and iVEC population to the fetal LEC and fetal VEC populations. In the PCA, we observed that the iLECs clustered with the fetal LECs and the iVECs clustered with the fetal VECs with a variance of 58.2% (PC1) between the LEC populations to the VEC populations. The 19.7% variance on PC2 may be ascribable to batch effects and technical variants (**Figure 5A**). Spearman correlation analysis with unsupervised clustering revealed that the iLECs clustered closely with fetal LECs from both the single cell RNA-seq datasets and the iVECs clustered closely with the fetal VECs. Notably, the degree of correlation between the iLECs and iVECs to their corresponding fetal cells was > 96% (**Figure 5B**).

**Figure 5.**
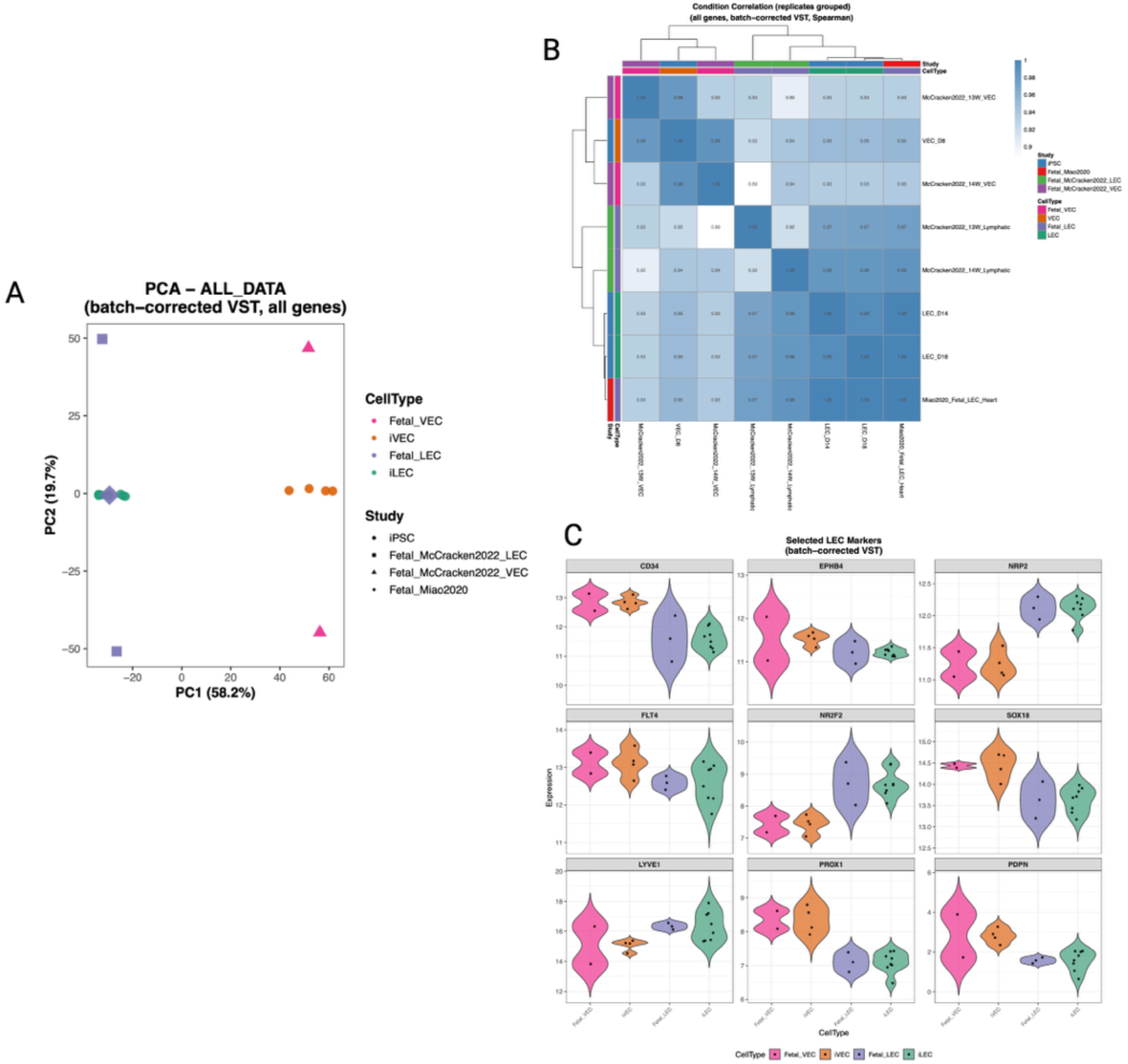
Comparison of iVECs and iLECs to fetal VECs and fetal LECs. **A.** PCA plot of iVECs and iLECs to fetal VECs and fetal LECs from heart endothelium of normal 13-week, 14-week and 12-week fetuses. **B.** Spearman correlation matrix comparing iVECs and iLECs to their respective fetal endothelial populations. **C.** Violin plot comparing gene expression of selected venous and lymphatic endothelial genes between iVECs and iLECs to each other and their respective fetal endothelial populations.

Finally, we compared expression of selected venous and lymphatic endothelial genes between the differentiated groups and the fetal groups. The expression levels depicted by violin plots further confirmed that the iLECs expressed selected genes at levels similar to those of fetal LECs, and the same was true for iVECs with respect to fetal VECs. As expected, the iVECs and fetal VECs had higher expression of *EPHB4*, *CD34, FLT4* and *PROX1,* while the iLECs and fetal LECs showed higher expression for *LYVE1* and *NR2F2* (**Figure 5C**). These data corroborated the previously shown mRNA expression (qPCR) and protein expression (flow cytometry).

Collectively these data supported the differentiation of iPSCs into robust VEC population with EGF and bFGF and LEC populations with EGF and VEGF-C, albeit at fetal stages.

## DISCUSSION

The advent of iPSC technology has enabled the generation of diverse cell types through differentiation protocols, providing researchers with human genetically matched cell models to advance our understanding of developmental and disease mechanisms. Continued development of such protocols will further these aims and expand the potential for personalized cell therapies in the future.^34^ In the context of lymphatic biology, despite the critical roles of the lymphatic vasculature in fluid homeostasis and immune cell trafficking and the dysregulation of which leads to lymphatic disease, the cellular mechanisms that drive lymphangiogenesis and lymphatic pathology remain relatively unexplored compared with their blood angiogenic counterparts. This is partly due to the scarcity of robust cell models to explore development and disease of the lymphatic system in a human context. In this present study, we developed a protocol that allowed us to differentiate iPSCs into LECs. It has been widely understood that lymphatic development begins with the trans-differentiation of VECs from the anterior cardinal vein.^10,35,36^ We, therefore, designed the protocol to recapitulate *in vivo* lymphatic development *in vitro* by directing venous endothelial to lymphatic endothelial transdifferentiation. Our goal was also to create an efficient feeder-free and serum-free protocol to circumvent the limitations of using feeder cells in differentiating protocols.^17,37^ Our protocol was successful in producing iVECs and iLECs that captured both venous and lymphatic marker expression as well as lymphatic functionality.

We demonstrated stepwise differentiation of the WTC11 human iPSC line, progressing sequentially through mesodermal, endothelial progenitor, and venous endothelial stages, followed by transdifferentiation to LECs. CHIR and bFGF were sufficient to induce primitive streak and mesodermal specification in the monolayer culture, after which we promoted endothelial progenitor fate using VEGFA and BMP4. We were consistently able to generate ∼24% CD31^+^/CD34^+^ endothelial progenitor population, which we then purified to ∼95% pure endothelial progenitor population. Following venous differentiation with EGF and bFGF, we found that mRNA levels of key venous endothelial markers, such as *EPHB4* and *NRP2,* were upregulated while key arterial endothelial markers were downregulated in our iVEC population (**Figure 2B**). Differences in expression levels of *NOTCH1* and *NRP2* were not significant but trended in the expected directions (lower *NOTCH1* and higher *NRP2* in iVECs compared to iAECs). Both *NOTCH1* and *NRP2* are crucial for angiogenesis, and while DLL4-mediated NOTCH1 activation is indicative of arterial specification, fetal NOTCH activity has been recorded in low VEGF venous conditions.^38–40^ Further, while *NRP2* expression is known to be biased to venous and lymphatic development, joint expression of *NRP1* and *NRP2* is shown in early angiogenesis and arterio-venous development.^41,42^ Our iVEC differentiation protocol was adapted from a previously published feeder-free and serum-free protocol, and expression profiles of our iVECs were consistent with findings from that study.^19^ It has also previously been suggested that VECs of the cardinal vein that express *FLT4* engage in VEGFR3/VEGF-C signaling and, in turn, are primed for lymphatic sprouting.^43,44^ Further, VECs that are primed to transdifferentiate and bud-off from the cardinal vein in lymph sacs express *PROX1*, the master lymphatic regulator^45^. Expression of *FLT4* and *PROX1* in our VEC population pointed towards a sub-population of the cells being primed for transdifferentiation and adopting a lymphatic phenotype (**Figure 2C**). We were, therefore, successful in creating a robust VEC population from endothelial progenitors, which we then directed towards lymphatic fate.

It is generally accepted that VEGF-C-mediated induction of the VEGR3 receptor is a crucial signaling pathway for lymphangiogenesis, and the sprouting of initial LECs from embryonic veins is attributed to VEGF-C/VEGFR-3 mediated signaling.^9,43,46^ Additionally, EGF (CCBE_1_) has been identified as a critical factor in regulating the activity of VEGF-C and promoting PROX1^+^ LEC budding from the cardinal vein.^47–49^ Considering this, we have shown that transdifferentiation of iVECs to iLECs with a combination of VEGF-C and EGF is sufficient to upregulate key lymphatic markers such as PROX1, PDPN, LYVE1 and NR2F2 while maintaining expression of SOX18 and FLT4. We also showed that our protocol generated a higher percentage of iLEC populations that expressed PROX1 and PDPN as compared to currently accepted OP9 feeder-reliant LEC differentiation protocols.^50,51^ On average across both differentiation timepoints, we obtained approximately 90% PROX1^+^ and 92% PDPN^+^ populations of iLECs. Moreover, we demonstrated that our iLECs were functionally similar to primary HDLECs in forming tubes as well are expressing key lymphatic proteins that could be stained for and imaged using immunocytochemistry.

Lastly, we compared the transcriptomic profiles from our iVECs and iLECs to those of human fetal VECs and fetal LECs. We found that gene expression in our iVECs matched well to fetal VECs and that gene expression of our iLECs was highly correct that of fetal LECs. We also observed similar expression levels of key venous and lymphatic markers between the differentiated cells and their fetal counterparts, further supporting the venous and lymphatic phenotypes of our differentiated cell populations.

In summary, our study describes an efficient feeder-free and serum-free approach to differentiate iPSCs into a robust lymphatic endothelial cell population. The protocol is designed to recapitulate key *in vivo* stages of lymphangiogenisis and, in doing so, enables reliable transdifferentiation of iPSC-derived VECs to LECs. By eliminating the need for feeder-cells, this approach improves scalability, reduces variability, technical complexity and experimental constraints commonly associated with feeder-cell dependent protocols. Given that our differentiated cell populations exhibit a fetal-like stage, they are especially well suited for modeling the molecular mechanisms that drive lymphatic development and disease. With this protocol, we hope to broaden access to LECs with a human-genetic background, and, in so doing, aid the progress of future discoveries in lymphatic biology and disease.

### Limitations of the study

Our transcriptomic analysis was limited to bulk RNA-sequencing analysis, which prevented us from parsing possible heterogeneity of both our venous and lymphatic populations. Further, our protocol relies on a 2D cell culture, which did not account for mechanical forces. The importance of mechanical forces in driving lymphatic development and maturity is well studied, and so future work can include examining the effects of mechanical stimuli and flow on transcriptional profile and signaling pathways in the differentiated iLECs.^52,53^

## METHODS

### Maintaining human iPSCs in culture

A wild-type human iPSC line, WTC11 (XY, mono-allelic ACTN2-mEGFP, obtained from Coriell Institute #AICS-0075-085), was grown, maintained and utilized in the experimental set up. The iPSCs were maintained in 6-well cell culture plates, where they adhered to a thin-layer of Matrigel (6.6% v/v, Corning), diluted in DMEM (Gibco) and supplemented with penicillin/streptomycin (5%, ThermoFisher). iPSC cultures were fed 2 ml of iPSC culture medium, which consisted of StemFlex medium (ThermoFisher) supplemented with penicillin/streptomycin (1%), daily. The cells were maintained in culture for roughly three days before being passaged when they reached around 80% confluency. iPSC cultures were dissociated with Accutase (Innovative Cell Technologies, Inc) to single cells and were replated at a 1:6 ratio on 6-well plates with the standard iPSC culture media which was supplemented with thiazovivin (2 µM, Selleckchem) for the first 24 h. The cell cultures were maintained in a 37 °C incubator with a normoxic environment (5% CO_2_).

### Monolayer differentiation of iPSCs into lymphatic endothelial-like cells

Preceding Day 0 of the differentiation (start of the differentiation), iPSCs were passaged and plated at a ratio of 1:12 on Matrigel-coated 24-well plates. The differentiation was split into two phases, which were primarily differentiated by the base media used. Phase 1 was adapted from a published protocol to differentiate iPSCs into venous endothelial cells^19^ and utilized RPMI 1640 (Gibco) that was supplemented with B27 minus insulin supplement (0.5x, Gibco), penicillin/streptomycin (1%), GlutaMAX (2 mM, Gibco), ascorbic acid (50 μg/mL, Sigma), apo-transferrin (150 μg/mL, R&D Systems), and monothioglycerol (50 μg/mL, Sigma). Day 0 of the differentiation was begun when the iPSC cultures were around 85% confluent and were introduced to Phase 1 media with CHIR-99021 (5 μM, Tocris) to induce the primitive streak. After 24 h, the media was changed to Phase 1 media supplemented with bFGF (50 ng/mL, R&D Systems). On Day 2, the media was changed to Phase 1 media supplemented with BMP4 (25 ng/ml, R&D Systems) and VEGF-A (50 ng/mL, R&D Systems) for 72 h. On Day 5, CD34^+^ cells were sorted out using MACS (Miltenyi, described below) and replated at 300,000 cells per well in a 24-well plate coated with Matrigel. These cells were cultured in Phase 1 media supplemented with EGF (10ng/ml, R&D Systems) and bFGF (20 ng/mL,) from Day 5 to Day 8 and was the media conditions for venous endothelial growth.

Phase 2 of the differentiation protocol aimed to transdifferentiate the venous endothelial cells to lymphatic endothelial-like cells. Media used for Phase 2 was the Endothelial Cell Growth medium MV 2 (#22121, PromoCell) supplemented with all cytokines present in the Supplement Mix (#22121, PromoCell), except for VEGF-165, and penicillin/streptomycin (1%). From Days 8-18, the media was changed from Phase 1 to Phase 2 media, which was additionally supplemented with EGF (10 ng/ml) and VEGF-C (50 ng/ml). The cells were passaged when they reached roughly 80% confluency to avoid spontaneous differentiation and cell death caused by overly confluent and crowded wells. The cells were passaged at a 1:6 ratio into 24-well plates. Day 18 was the end point of the differentiation. Throughout the entire differentiation, the cells were maintained at 37 °C in a hypoxic incubator (5% CO_2_, 5% O_2_, 90% N_2_), and the media was changed every other day.

### Cell sorting and flow cytometry

For analysis of the purified endothelial progenitor population on Day 5, cells were dissociated using collagenase type II (0.6 mg/ml, Worthington) in HBSS (Corning) supplemented with HEPES (5 mM, Sigma) and incubated at 37 °C for 1 h. Post dissociation, the cells were collected in wash media consisting of DMEM (Gibco), bovine serum albumin (BSA, 0.05%, Gibco), DNAse (10 mg/mL, Millipore) and thiazovivin (2 µM), where they were passed through a 40-μm cell strainer (Fisher Scientific) and collected on ice. MACS was used to enrich the cell population to a purity of roughly 95% of CD34^+^/CD31^+^ double-positive endothelial cells. Briefly, the dissociated cells were incubated with human anti-CD34 microbeads (Miltenyi #130-046-702) in EasySep Buffer (Stem Cell Technologies) for 15 min on ice, in accordance with the method described in the instruction manual of the kit. The antibody-stained cells were then purified using an LS column, and a minimum of three washes through the column using the EasySep Buffer were performed to increase purity of the sorted cells.

For flow cytometry analysis, the enriched CD34^+^ cell population obtained on Day 5, was then resuspended in FACS staining buffer consisting of PBS^-/-^ (Gibco) supplemented with FBS (10%, Gibco), DNase (10 mg/mL, Millipore), and thiazovivin (2 µM) and labeled with anti-CD31 APC (1:100, Invitrogen) and anti-CD34 PE-Cy7 (1:100, BioLegend) for 1 h on ice. After an hour of incubation, the cells were washed three times using PBS^-/-^ in excess, after which they were resuspended in FACS running buffer consisting of PBS^-/-^ supplemented with FBS (1%), penicillin/streptomycin (2%), DNase (10 mg/mL), thiazovivin (2 µM), and DAPI (0.1 mg/mL, Sigma) and analyzed using flow cytometry.

For flow cytometry analysis on Day 14 and 18, cells were dissociated using Accutase for 15 min and collected in two 15-ml centrifuge tubes. Cells in tube 1 were stained with anti-PDPN PE (1:100, BioLegend) and anti-CD34 PE-Cy7 (1:100, BioLegend) using the same staining and FACS buffer steps as mentioned above. Cells in tube 2 were fixed with 200 ul of 4% PFA in PBS (ThermoFisher) for 10 min at room temperature. The cells were then washed three times using PBS followed by permeabilization with 0.1% Triton X-100 in PBS for 10 min. The cells were then washed with PBS again before being resuspended in the FACS staining buffer (described above) and stained with anti-CD34 PE-Cy7 (1:100, BioLegend) and anti-PROX1 PE (1:100, NovusBio). The stained cells were then resuspended in FACS running buffer (described above) and analyzed using flow cytometry.

### In-vitro Matrigel tube formation assay

On Day 18 of the protocol, cells growing on four wells of a 24-well plate were serum starved using DMEM and 0.1% FBS for 24 h. After 24 h, the cells were dissociated with Accutase to conduct a tube formation assay, which was adapted from a previously published protocols.^54,55^ 2 × 10^4^ differentiated cells suspended in the Phase 2 media were then plated on µ-slide 8-well high glass bottom plates (#80807, Ibidi) that were coated with 150 µl of growth-factor reduced Matrigel (6.6% v/v). The Matrigel had a thickness of 1.5 mm. Tube formation was observed at 4 h, and the tubes were fixed using 4% PFA for 10 min to prepare for immunofluorescence.

### Immunocytochemistry

On Day 14 of the protocol, cells from two wells of the 24-well plate growing in the lymphatic endothelial condition (Phase 2 media) were dissociated using Accutase and replated on 8-chamber glass bottom μ-slides (Ibidi) coated with Matrigel (6.6% v/v). The replated cells were maintained in Phase 2 media supplemented with EGF (10 ng/ml) and VEGF-C (50 ng/ml) in the 37 °C hypoxic incubator (5% CO_2_, 5% O_2_, 90% N_2_). On Day 18, the cells maintained on the 8-chamber μ-slides were fixed with 4% PFA in PBS for 10 min at room temperature, followed by permeabilization with 0.1% Triton X-100 in PBS for 10 min. The cells were then washed with PBS and blocked for 1 h at room temperature with PBS containing BSA (1%), glycine (22.52 mg/mL, BioRad), Tween-20 (0.1%, Sigma), and sodium azide (0.02%, Sigma). They were then labeled with primary antibodies and incubated at 4 °C for 24 h in PBS containing BSA (1%), Tween-20 (0.1%), and sodium azide (0.02%) after which they were washed with PBS three-times for 10 min each and stained with secondary antibodies for 1 h at room temperature, followed by another wash with PBS. Cell nuclei were stained with DAPI (0.25 mg/mL, Sigma) in PBS for 10 min at room temperature. The following primary antibodies were used: anti-PROX1 (1:100, Proteintech), anti-CD31 (1:100, Invitrogen), anti-LYVE1 (1:50, Invitrogen), anti-VECAD (1:100, Cell Signaling Technology), and anti-PDPN (1:100, BioLegend). The secondary antibodies used were goat anti-rabbit IgG Alexa Fluor 488 (1:400, Invitrogen) and goat anti-mouse IgG Alexa Flour 568 (1:400, Invitrogen). The stained cells were then washed with PBS once, after which 200 μL of PBS^-/-^ was added to each well to prevent the cells from drying out. Stained cells were imaged on an Andor Dragonfly 620 spinning disk microscope. Additionally, tubes fixed on Day 19 were permeabilized, blocked and stained using the protocol described above. Single cells as well as tubes were imaged to detect fluorescence of the primary antibodies used. Images were analyzed using FIJI software.

### Quantitative reverse transcriptase PCR

Cells were dissociated on Days 8, 14 and 18 respectively, and total RNA from the samples was isolated using the Qiagen RNeasy Mini Kit. Cells were collected in 1.5-ml RNAse-free microfuge tubes and lysed with Buffer RLT containing *β*-mercaptoethanol (Sigma) per the manufacturer’s instructions. The samples were then run through a RNeasy column for RNA isolation. Both Part 1 and 2 of RNeasy Mini Kit protocol were followed. In accordance with the instruction manual from the manufacturer, DNase treatment was performed on-column and total RNA was eluted in RNase-free H_2_O. The quality of the RNA was determined with the Agilent 2100 Bioanalyzer and the RNA 6000 Nano chip, where the RIN numbers were confirmed to be > 8.0 for all samples. The SuperScript IV VILO Master Mix (Invitrogen) was used to convert the total RNA to cDNA, and qRT-PCR was performed on a ViiA 7 Real-Time PCR System (Applied Biosystems) using the PowerTrack SYBR Green Master Mix (ThermoFisher). All experiments were run in triplicates using the ΔΔCt method relative to a human reference cDNA (Takara) and the housekeeping gene TBP. RNA samples from Day 8 were from iVECs and iAECs and were analyzed separately from RNA samples collected from Day 14 and Day 18 cells. Primer sequences are listed in Table S1.

### Bulk RNA sequencing analysis

Cells on Day 8, 14 and 18 were collected and lysed with Buffer RLT containing *β*-mercaptoethanol. Total RNA extraction as well as library prep and sequencing were performed by Novogene. All total RNA samples were confirmed to have a RIN > 8.0. Reads were aligned by Novogene to the human reference genome (hg38), and downstream analyses were initiated using vendor-delivered BAM files. Gene-level counts were generated using featureCounts(v2.18.0)^56^ with the supplied GTF annotation (exon-level assignment; summarized by gene_id). Gene symbols were appended from the reference annotation, and exon-based gene lengths were computed for TPM estimation. Count matrices were analyzed in R with DESeq2(v1.44.0).^30^ Genes with low counts (sum < 10 across all samples) were filtered prior to analysis. Differential expression was modeled with replicate as a batch term and group as the biological factor (design ∼ batch + group). Size-factor normalization and variance-stabilizing transformation were performed in DESeq2, and batch-corrected transformed values for visualization were obtained using limma(v3.60.6).^31^ Exploratory analyses included PCA, UMAP, multidimensional scaling using 1 − Spearman correlation as a distance metric, sample-to-sample correlation analyses, and heatmaps of selected marker genes (row-scaled variance-stabilized expression). Differential expression was performed in DESeq2 (Wald test), and log2 fold changes were shrinkage-estimated using ashr (v2.2.63).^57^ Gene set enrichment analysis was performed with fgsea (v1.30.0)^58^ using MSigDB gene sets (v10.0.1).^59^

For comparison with fetal endothelial populations, published fetal scRNA-seq datasets from Cao et al.,^59^ Miao et al.,^33^ and McCracken et al.^60^ were downloaded and processed in R. Raw UMI matrices were subset to annotated endothelial populations and converted to pseudobulk profiles by summing counts across biologically matched groups (e.g., organ or developmental stage). Gene identifiers were harmonized using Ensembl IDs (version suffixes removed; duplicate mappings collapsed by summation), and analyses were restricted to genes detected in both bulk and pseudobulk datasets. The combined bulk–fetal matrix was normalized in DESeq2 using sfType = “poscounts” followed by variance-stabilizing transformation; study and replicate-associated effects were removed using limma, and concordance was assessed by PCA, hierarchical clustering, marker-gene heatmaps, and Spearman correlation. Raw data from iPSC-derived populations can be found at GSE324673.

### Statistical analysis

GraphPad Prism Software (GraphPad Software639 Inc., La Jolla, CA) was used to perform all statistical analysis. All data reporting statistics were performed with a minimum of three replicates. Paired *t*-test analysis was used to compare between two different cell types, as well as two different type points respectively. All results report a mean ± SEM, along with individual datapoints shown. Significance levels are reported as: *p < 0.5, **p < 0.01, ***p < 0.001, ****p < 0.0001.

## Acknowledgments

We gratefully acknowledge the Icahn School of Medicine at Mount Sinai, particularly the Department of Stem Cell Biology and Regenerative Medicine and the Developmental, Regenerative, and Stem Cell training area within the Graduate School of Biomedical Sciences for institutional support. We also extend our sincere thanks to Dr. Nicole Dubois for her thoughtful feedback and unwavering support as well as Dr. Sushmita Sahoo for her technical support which was crucial in developing the tube formation assay. We acknowledge the Flow Cytometry Core and the Microscopy Core for essential technical resources and assistance. This work was supported by the National Institutes of Health (R01 HL173006). Finally, the differentiation schematic was created with BioRender.com.

## Author contributions

Conceptualization, A.P., C.Z.L and B.D.G.; methodology, A.P and B.D.G.; validation: A.P., S.N. and B.D.G.; formal analysis, A.P.; S.P and B.D.G.; investigation, A.P; resources, B.D.G.; data curation: A.P., S.P.; writing – original draft: A.P. and B.D.G.; visualization: A.P. and S.P.; supervision: B.D.G.; project administration: B.D.G.; funding acquisition: B.D.G.

## Declaration of Interest

The authors declare no competing interests.

## Data and materials availability

Additional information and data contributing to the study can be found in **Supplemental information**.

## Supplemental information

**Figure S1:**
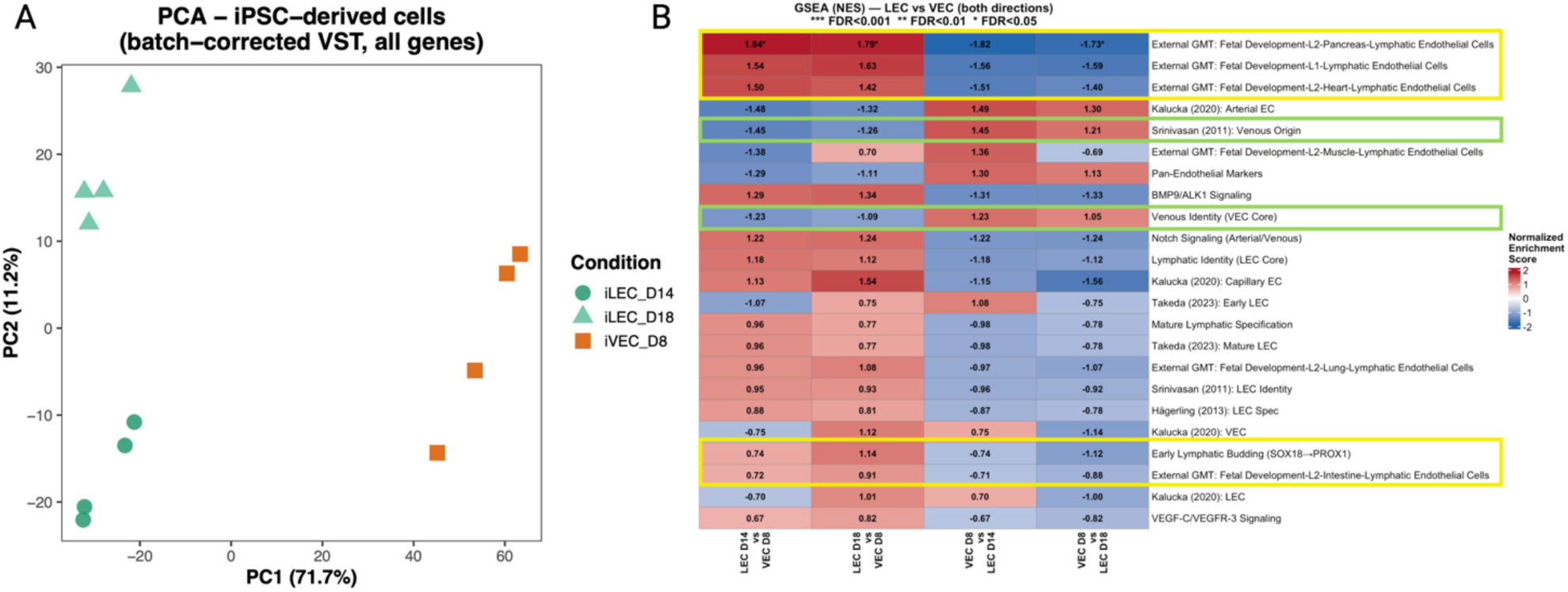
Bulk-RNA sequencing analysis of iVECs and iLECs on Day 8, 14 and 18, related to Figure 5. **A.** PCA plot of iVECs and iLECs at Day 8, 14 and 18. **B.** GSEA heatmap for enriched genes and pathways in iVECs and iLECs (on Day 14 and 18).

**Table S1.**
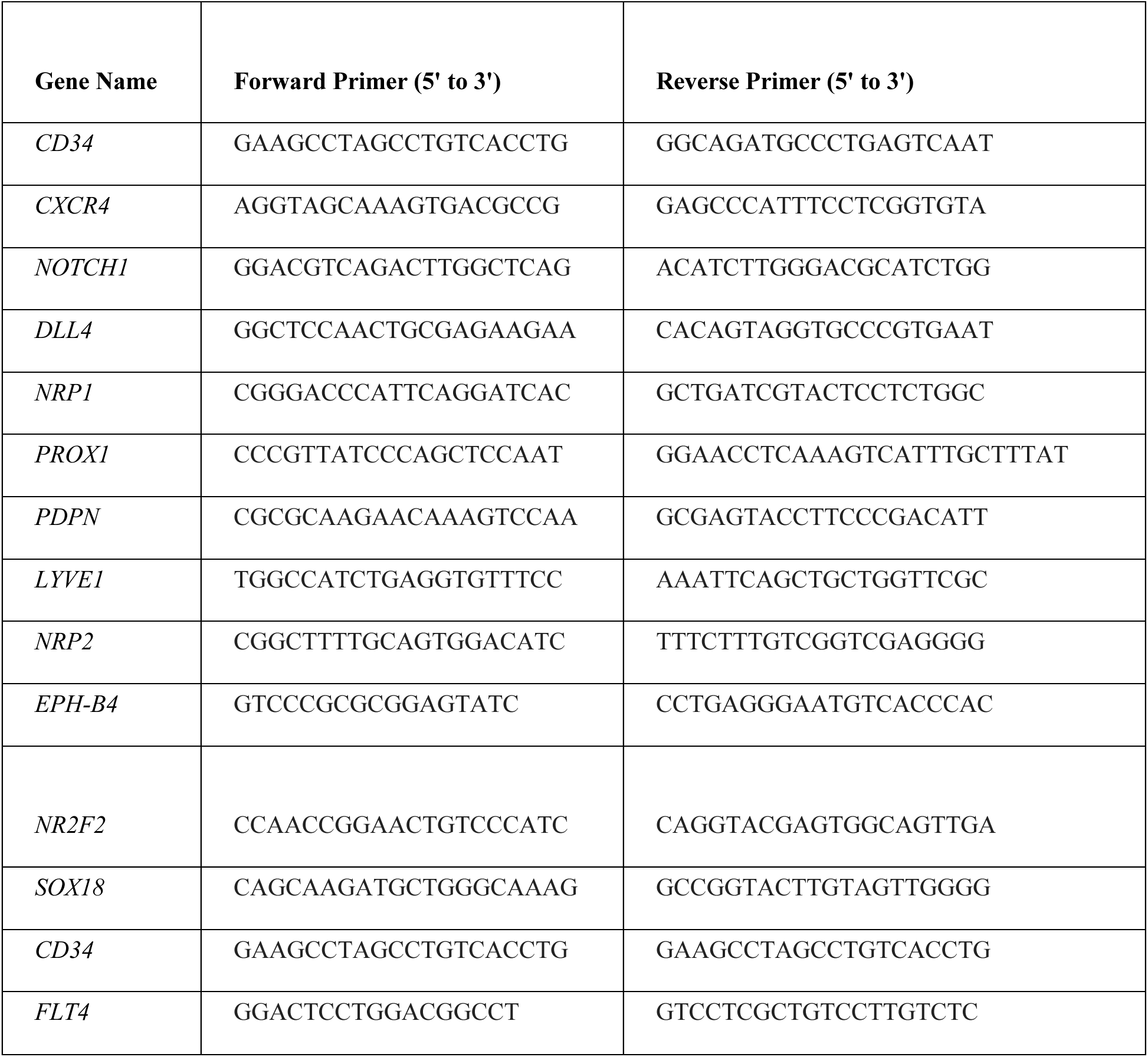
List of Primers, related to Methods.

## Notes

### Competing Interest Statement

The authors have declared no competing interest.

