## Supplementary material for "Feeder-Free Generation of Lymphatic Endothelial Cells from Human Induced Pluripotent Stem Cells": Supplementary information only.docx

**Supplemental information**

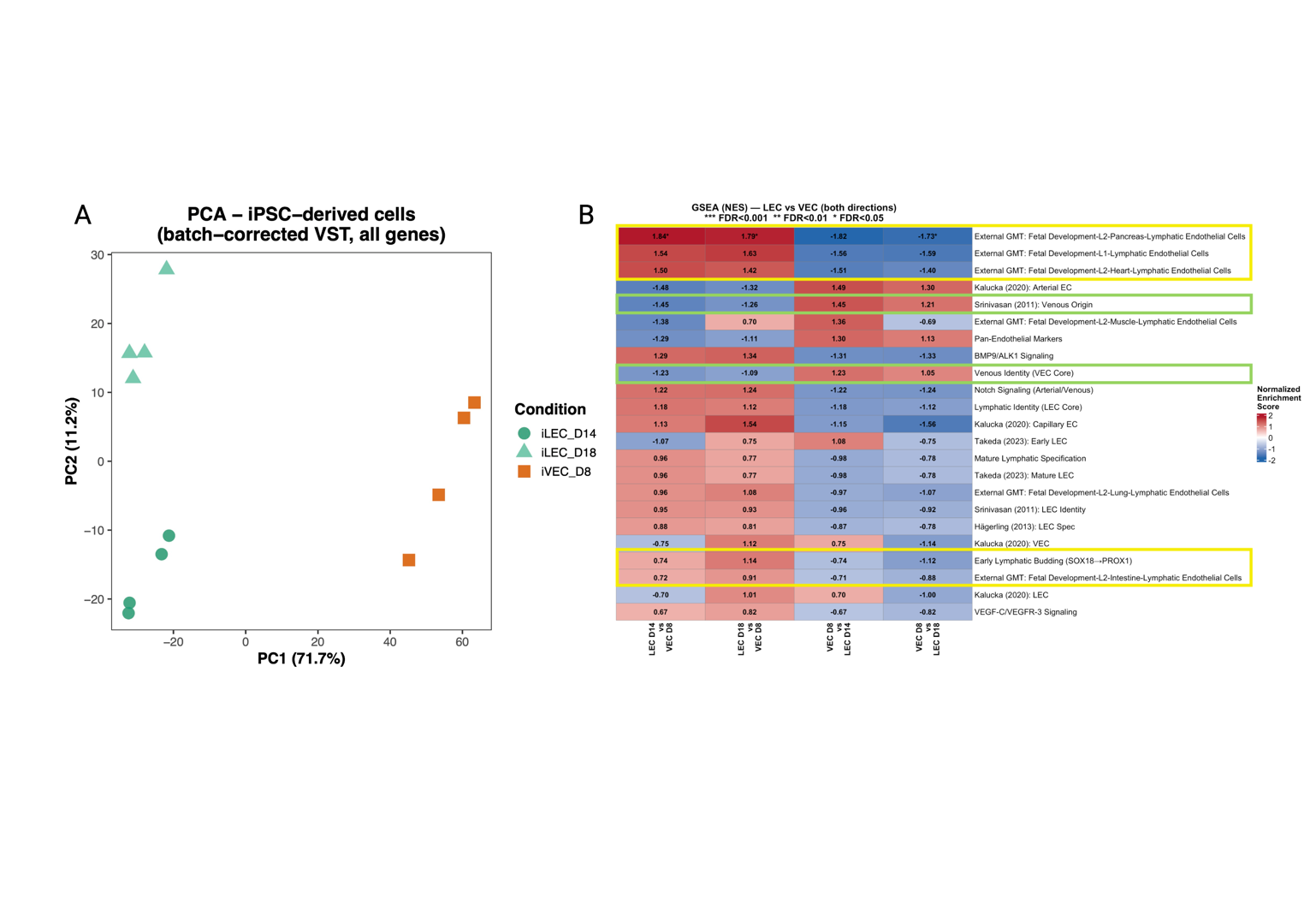

**Figure S1:** Bulk-RNA sequencing analysis of iVECs and iLECs on Day 8, 14 and 18, related to Figure 5. **A.** PCA plot of iVECs and iLECs at Day 8, 14 and 18. **B.** GSEA heatmap for enriched genes and pathways in iVECs and iLECs (on Day 14 and 18).

**Table S1 – List of Primers, related to Methods**

| **Gene Name** | **Forward Primer (5' to 3')** | **Reverse Primer (5' to 3')** |
| --- | --- | --- |
| *CD34* | GAAGCCTAGCCTGTCACCTG | GGCAGATGCCCTGAGTCAAT |
| *CXCR4* | AGGTAGCAAAGTGACGCCG | GAGCCCATTTCCTCGGTGTA |
| *NOTCH1* | GGACGTCAGACTTGGCTCAG | ACATCTTGGGACGCATCTGG |
| *DLL4* | GGCTCCAACTGCGAGAAGAA | CACAGTAGGTGCCCGTGAAT |
| *NRP1* | CGGGACCCATTCAGGATCAC | GCTGATCGTACTCCTCTGGC |
| *PROX1* | CCCGTTATCCCAGCTCCAAT | GGAACCTCAAAGTCATTTGCTTTAT |
| *PDPN* | CGCGCAAGAACAAAGTCCAA | GCGAGTACCTTCCCGACATT |
| *LYVE1* | TGGCCATCTGAGGTGTTTCC | AAATTCAGCTGCTGGTTCGC |
| *NRP2* | CGGCTTTTGCAGTGGACATC | TTTCTTTGTCGGTCGAGGGG |
| *EPH-B4* | GTCCCGCGCGGAGTATC | CCTGAGGGAATGTCACCCAC |
| *NR2F2* | CCAACCGGAACTGTCCCATC | CAGGTACGAGTGGCAGTTGA |
| *SOX18* | CAGCAAGATGCTGGGCAAAG | GCCGGTACTTGTAGTTGGGG |
| *CD34* | GAAGCCTAGCCTGTCACCTG | GAAGCCTAGCCTGTCACCTG |
| *FLT4* | GGACTCCTGGACGGCCT | GTCCTCGCTGTCCTTGTCTC |
